# MPL36, a major plasminogen (PLG) receptor in pathogenic *Leptospira,* has an essential role during infection

**DOI:** 10.1101/2023.03.24.534066

**Authors:** Weinan Zhu, Felipe J. Passalia, Camila Hamond, Cecília M. Abe, Albert I. Ko, Angela S. Barbosa, Elsio A. Wunder

**Affiliations:** Department of Epidemiology of Microbial Diseases, Yale School of Public Health, New Haven, CT, United States; Laboratory of Vaccine Development, Instituto Butantan, São Paulo, Brazil; Laboratory of Bacteriology, Instituto Butantan, São Paulo, Brazil; Gonçalo Moniz Institute, Oswaldo Cruz Foundation; Brazilian Ministry of Health; Salvador, Brazil

## Abstract

Leptospirosis, a zoonosis with worldwide distribution, is caused by pathogenic spirochetes belonging to the genus *Leptospira*. Bacterial outer membrane proteins (OMPs), particularly those with surface-exposed regions, play crucial roles in pathogen dissemination and virulence mechanisms. Here we characterized the leptospiral Membrane Protein L36 (MPL36), a rare lipoprotein A (RlpA) homolog with a C-terminal Sporulation related (SPOR) domain, as an important virulence factor in pathogenic *Leptospira*. Our results showed that MPL36 is surface exposed and expressed during infection. Recombinant MPL36 (rMPL36) showed high plasminogen (PLG)-binding ability determined by lysine residues of the C-terminal region of the protein, with ability to convert bound-PLG to active plasmin. Using Koch’s molecular postulates, we determined that a mutant of *mpl36* has a reduced PLG-binding ability, leading to a decreased capacity to adhere and translocate MDCK cell monolayers. Using recombinant protein and mutant strains, we determined that the MPL36-bound plasmin (PLA) can degrade fibrinogen. Finally, our *mpl36* mutant had a significant attenuated phenotype in the hamster model for acute leptospirosis. Our data indicates that MPL36 is the major PLG binding protein in pathogenic *Leptospira*, and crucial to the pathogen’s ability to attach and interact with host tissues during infection. The MPL36 characterization contributes to the expanding field of bacterial pathogens that explore PLG for their virulence, advancing the goal to close the knowledge gap regarding leptospiral pathogenesis while offering a novel potential candidate to improve diagnostic and prevention of this important zoonotic neglected disease.

**Author Summary:** As part of their diverse virulence machinery, bacterial pathogens bind to human plasminogen (PLG) providing them with a proteolytic platform that promotes invasiveness, dissemination, and virulence. Leptospirosis is the leading zoonotic disease in morbidity and mortality worldwide.

The burden of this neglected disease will continue to raise given the effects of climate change and social inequality, important drivers of disease. Furthermore, the gap of knowledge regarding leptospiral pathogenesis has negatively impacted the development of sensitive diagnostic tools and effective prevention methods. Previous studies have shown that pathogenic *Leptospira*, the causative agent of leptospirosis, can interact with PLG through different protein candidates. In this work, we characterized one of those candidates, Membrane Protein L36 (MPL36), as the main leptospiral plasminogen binding protein. Using genetically modified mutants, *in vivo*, and *in vitro* assays we provided evidence that MPL36 can bound PLG, promotes adherence to host cells and subsequent translocation, and degrades fibrinogen by converting bound-PLG to PLA, thus essential to leptospiral virulence. This work contributes to the growing field of bacterial pathogens exploring PLG to increase their virulence, while highlighting important new knowledge on leptospiral pathogenesis. MPL36 is an important candidate to be explored on the continued effort to improve diagnostic and prevention of this important zoonotic disease.

## Introduction

Interaction with the human plasminogen (PLG) system significantly contributes to the virulence of many bacterial pathogens by equipping them with a proteolytic platform that enables bacterial invasiveness and tissue destruction [1]. Sequestration of PLG with further activation into plasmin (PLA) is crucial for bacterial survival in the host environment, since surface-bound PLA degrades fibrin clots, ECM molecules, and host’s innate immune proteins facilitating dissemination and escape from immune responses [2, 3]. The pathogenic spirochete *Leptospira,* the causal agent of the life-threatening infectious disease leptospirosis, is known to interact with host’s fibrinolytic system to ensure dissemination during the infection process [4].

Leptospirosis is a zoonotic disease of worldwide occurrence, which has a significant public health impact especially in low-income tropical and sub-tropical countries [5]. Globally, more than one million cases and approximately 60,000 deaths from leptospirosis are reported each year [6]. The severe form of leptospirosis, accounting for 10% of all cases, may be fatal due to bleeding manifestations and acute kidney injury. Mortality rates of up to 74% have been reported in patients who developed leptospirosis-associated pulmonary hemorrhage-syndrome [7–12]. The disease also affects the agricultural industry, causing abortions, infertility, and death in livestock [5, 13]. Since there are no preventive measures to control the infection in humans, leptospirosis remains a threat in developing countries lacking proper sanitation systems [6, 13, 14].

Currently classified into 69 species, the genus *Leptospira* was recently grouped in two major clades, namely the “Saprophytes” composed of free-living, nonpathogenic species, and the “Pathogens” which comprise species known to cause disease in humans and animals (P1) or species whose virulence status awaits confirmation, formerly called “Intermediates” (P2) [15, 16]. Pathogenic *Leptospira* enter the host through injured skin or mucous membranes, and rapidly reach the bloodstream [17, 18], due to their efficient swimming and crawling motilities through viscous environments [19, 20]. The first step in the process of leptospiral infection is cellular adhesion, mediated by surface proteins interacting with various components of the extracellular matrix (ECM) [21]. Several putative adhesins that bind host ECM proteins have been identified in pathogenic *Leptospira* [22], but consistent evidence regarding cell/ECM-binding activity was only demonstrated for a few of them, such as the Leptospiral immunoglobulin-like proteins A (LigA) and B (LigB) [23] and the outer membrane protein L1 (OmpL1) [24]. These spirochetes are also equipped with additional mechanisms for host colonization, including the secretion of proteases that display proteolytic activity against ECM and plasma proteins [25], and the subversion of host proteases such as PLG through surface receptors [26, 27]. Conversion of bound PLG into PLA by specific activators generates a proteolytic platform on the leptospiral surface supposedly increasing its invasiveness potential.

Several leptospiral proteins were previously described to act as PLG receptors. Some of them are well studied outer membrane proteins (OMPs) such as endostatin-like protein A (LenA), Leptospiral immunoglobulin-like proteins A (LigA) and B (LigB) and LipL32 [26, 28, 29]. Others are moonlighting proteins among which the elongation factor-thermal unstable (Ef- Tu) [30] and the metabolic enzyme enolase [31], also known to act as PLG receptors in other bacteria. The interaction of other less studied leptospiral surface proteins with PLG was also reported [28, 32]. Of special interest within this group is the Membrane Protein L36 (MPL36) encoded in *Leptospira interrogans* strain Fiocruz L1-130 by the gene *lic10054*. MPL36 was shown to bind human PLG with high affinity, presenting the lowest dissociation constant (*K*D) value among all proteins tested [32]. In the current study, using a random mutant [33] that lacked expression of MPL36, we performed genomic manipulation, *in vivo* studies and *in vitro* assays to demonstrate and characterize the mechanism in which pathogenic *Leptospira*, by engaging MPL36, successfully exploits host PLG for its dissemination within the host, critical for establishing infection.

## Results

### Disruption of *mpl36* does not affect cell morphology and growth rate

MPL36 is a Rare lipoprotein A (RlpA) homolog, initially described in *Escherichia coli* [34, 35], and has a double-psi beta-barrel domain (DPBB) and a C-terminal Sporulation related (SPOR) domain [36] (Fig 1A). RlpA is localized at the septal ring in *E. coli*, but a precise role for this protein has not yet been defined in this organism, since no obvious phenotype associated with cell division was observed in *rlpA* mutants [34]. In *Pseudomonas aeruginosa*, RlpA was characterized as a peptidoglycan hydrolase digesting “naked” glycans, and the protein was shown to be crucial for proper separation of daughter cells and maintenance of rod morphology [37, 38]. Based on these facts, we assessed cell morphology and growth ability of our Manilae strains. No differences in cell morphology or motility were observed while comparing the Δ*mpl36* mutant, the wide-type, and the complemented strains (data not shown). Similarly, comparable growth rates in EMJH medium at 30°C were exhibited by all three strains (S1C Fig), indicating that the disruption of the *mpl36* gene does not affect the ability of the strain to multiply and grow *in vitro*.

**Fig 1.**
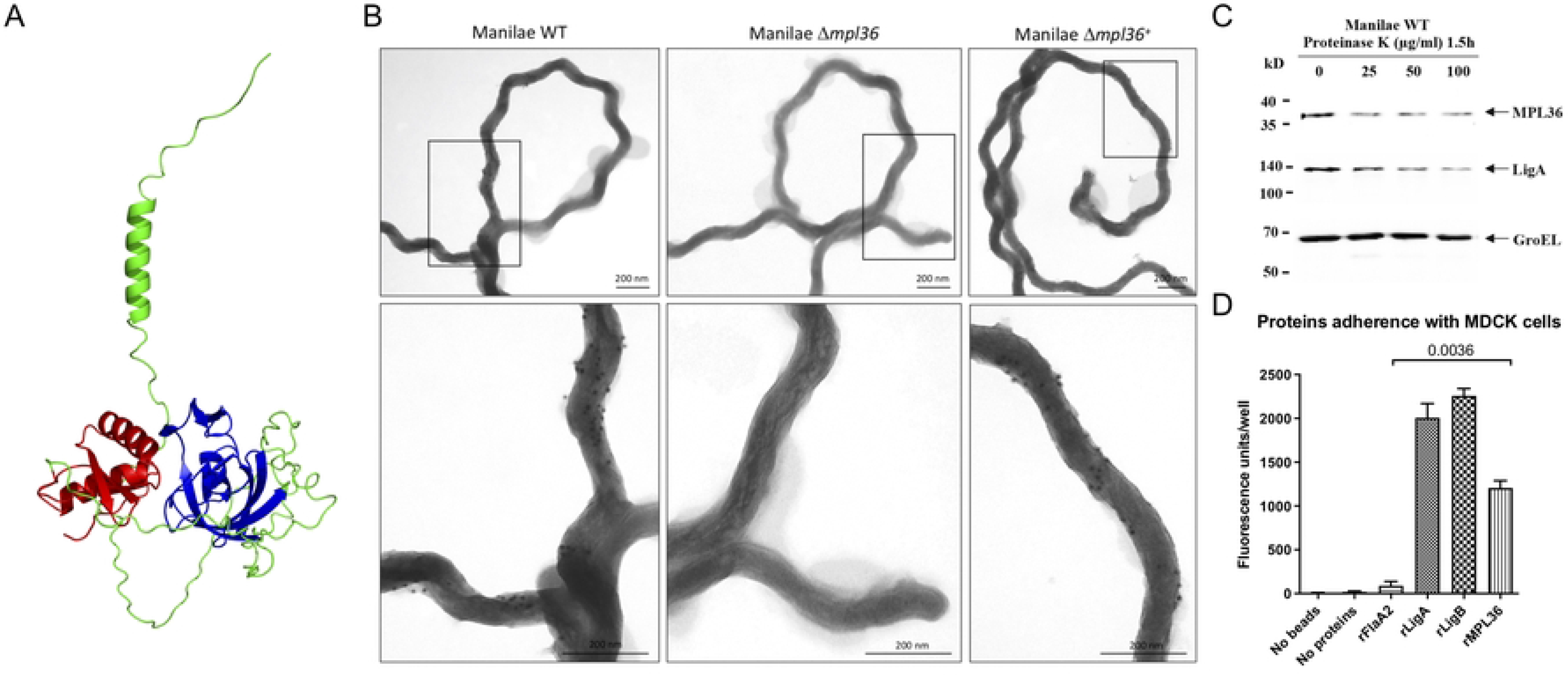
Structure and surface localization of MPL36 in *L. interrogans* serovar Manilae L495 and rMPL36 interaction with host epithelial cells. (A) Predicted model of *L. interrogans* MPL36 with the double-psi beta-barrel domain (DPBB) (blue), a conserved region of RlpA proteins (green), and the SPOR domain (red) in the C-terminal. (B) Immunogold labeling of WT, Δ*mpl36*, and Δ*mpl36^+^ Leptospira* strains were performed using polyclonal rabbit antiserum against MPL36 and goat anti-rabbit IgG labelled with 10 nm colloidal gold particles. Cells were visualized using 2% UA negative staining. (C) Whole intact spirochetes were incubated with different concentrations of Proteinase K (25-100 μg/mL), and western-blot analysis was conducted using polyclonal rabbit antisera against MPL36, LigA (positive control), and GroEL (negative control). (D) Recombinant proteins coated with fluorescent latex beads were incubated with immobilized MDCK cells and this interaction was assessed by fluorescent emission. LigA and LigB were used as positive control, while FlaA2 was used as negative control. The results are represented as mean ± standard deviation of two independent experiments.

### Complementation of the *mpl36* gene restores expression of the protein

*L. interrogans* serovar Manilae Δ*mpl36* mutant strain was generated by Himar1 transposon mutagenesis, with the transposon insertion on position 921,626 of the Manilae genome (558 bp from the start codon of the gene) (S1A Fig). The complemented strain was generated by the insertion of the transposon carrying the gene *mpl36* and a spectinomycin resistance cassette. After semi-PCR and sequencing screening, we identified four complemented strains, all of which exhibited the transposon in an intergenic region. The complemented strain Manilae Δ*mpl36*^+^, with the complementing construct inserted at chromosome position 292,919, was chosen for this study (S1A Fig). PCR and Sanger sequencing confirmed the presence of the *mpl36* gene in the WT and complemented strains (data not shown).

Immunoblotting of bacterial whole-cell lysates with a rabbit polyclonal antibody raised against rMPL36 allowed detection of the native protein in both the WT and complemented (Δ*mpl36^+^*) strains. A specific protein band with the apparent molecular mass of ∼40 kDa was not visible in the mutant (Δ*mpl36*) strain (S1B Fig). This result confirmed the disruption of *mpl36* gene, and consequently, lack of expression of MPL36 in the mutant, as well as restoration of protein production by the complemented strain.

### MPL36 is a surfaced exposed *Leptospira* protein that binds to host cells and PLG

MPL36 was predicted to be an outer membrane lipoprotein according to *in silico* analysis using lipoP and TMHMM software. Immunogold labeling using the WT, Δ*mpl36*, and Δ*mpl36^+^* strains demonstrated that anti-MPL36 antibodies specifically labeled the surface of the WT and Δ*mpl36^+^* strains and did not bind to the surface of the Δ*mpl36* strain (Fig 1B). Surface exposure was further assessed by proteinase K treatment of intact leptospires (*L. interrogans* serovar Manilae L495). Both MPL36 and the well-characterized surface protein LigA [5, 23] were sensitive to proteinase K-mediated degradation while the cytoplasmic control protein GroEL was not, providing additional evidence for MPL36 cell surface localization (Fig 1C). Immunoblot assay on the phase-partitioned fractions of *Leptospira* detected the MPL36 and surface LigA protein in the hydrophobic detergent phase, while the cytoplasmic protein GroEL [39] was portioned into the aqueous phase, and the periplasmic protein FlaB [26] was detected in the protoplasmic cylinder fraction (S2A Fig). Taken together, we demonstrated here the subcellular localization of MPL36 in *Leptospira* as an outer membrane protein.

The role of rMPL36 in adhesion to mammalian cells and ECM components was further investigated. Fluorescent latex beads coated with rMPL36 bound to MDCK cells whereas uncoated beads or those coated with rFlaA2 (negative control) did not display cell-binding activity (Fig 1D). As expected, rLigA and rLigB (positive controls) exhibited significant binding to MDCK cells (Fig 1D).

The interaction of rMPL36 with ECM proteins and PLG was also evaluated. No significant binding to fibronectin (*p* > 0.9, Fig 2A) or laminin (*p* = 0.9, Fig 2B) was detected with rMPL36, unlike what was observed with rLigA and rLigB proteins (positive controls). However, rMPL36 displayed a significant binding capacity to human PLG compared to all other recombinant proteins used either as positive or negative controls (*p* < 0.0001, Fig 2C).

**Fig 2.**
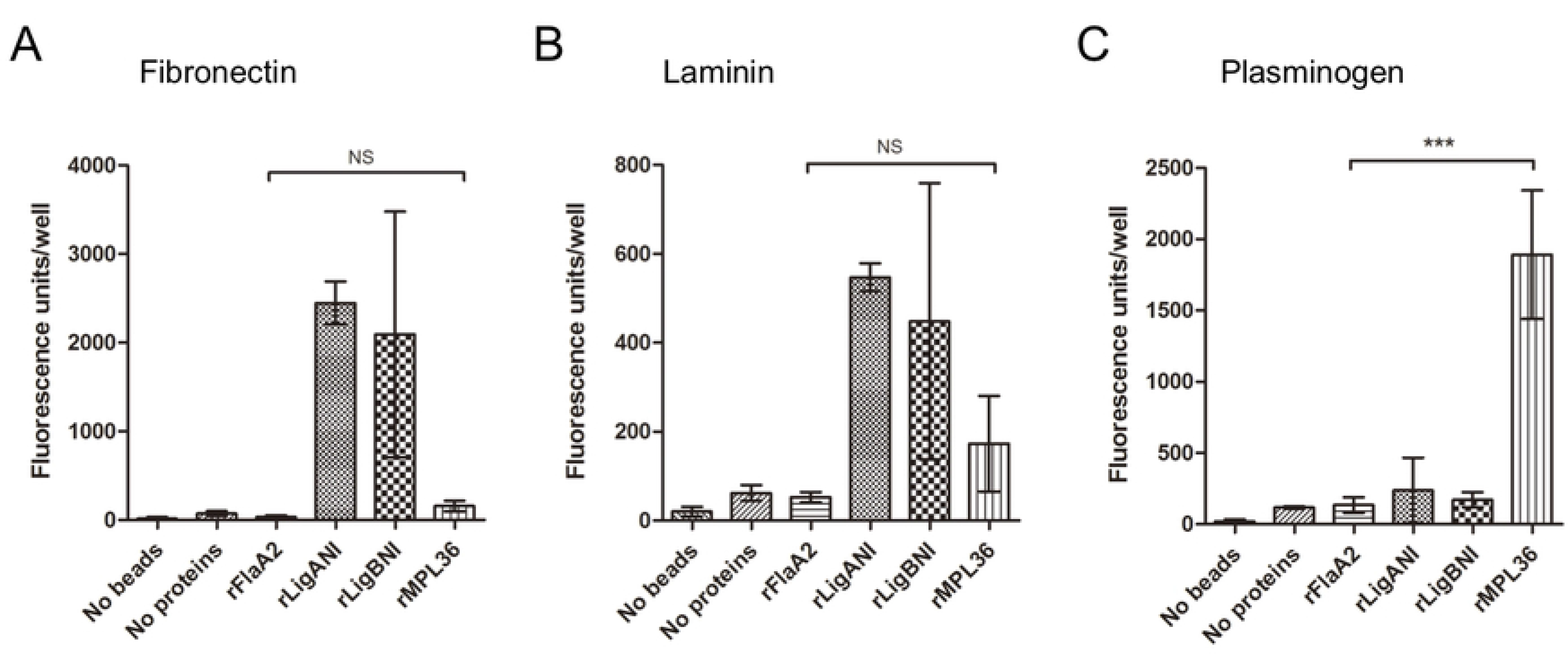
**Binding of rMPL36 to ECM components and plasminogen (PLG)**. 96-well plates were coated with 1 µg of fibronectin (A), laminin (B), and PLG (C). 0.2 nmol of the recombinant proteins MPL36, LigA, LigB or FlaA2 coated with fluorescent latex beads were added per well, and the binding was assessed by fluorescent emission. A control using uncoated fluorescent latex beads was used to measure background signal. Data represent the mean ± standard deviation of results from three independent experiments (\*\*\**p* < 0.0001).

Furthermore, rMPL36 is recognized by sera from individuals with laboratory confirmed severe leptospirosis from Salvador, Brazil. Convalescent sera from those individuals have a high level of antibodies against MPL36 in both IgM (S2B Fig) and IgG (S2C Fig) when compared to sera from healthy individuals (*p* = 0.0023 and *p* < 0.001). No statically significant difference was observed in the acute sera (S2B and C Fig), although 48% of these sera had IgM antibodies levels higher than the identified threshold (S2B Fig, *p* = 0.08), further confirming its expression during host infection and suggesting a potential role on the initial phases of the disease.

Taken together, those results suggest that rMPL36 can directly associate with MDCK cells, which is consistent with its surfaced-exposed localization and expression in the host during leptospirosis, in addition to its ability to bind to PLG, as previously described [28].

### MPL36 binds PLG by its C-terminal lysine residues and MPL36-bound PLG is converted to PLA

To be proteolytically active, PLG needs to be converted in its active form, PLA, by urokinase or tissue PLG activators (uPA or tPA, respectively). The ability of rMPL36-bound PLG to be converted into PLA was then assessed. By exogenously supplying uPA to rMPL36- bound PLG, the newly generated PLA was able to significantly cleave the chromogenic substrate D-valyl-leucyl-lysine-ρ-nitroanilide dihydrochloride compared to BSA control (p < 0.001) (Fig 3A). Significant PLA activity was not detected with rLigA, rLigB, or rFlaA2 (Fig 3A). By ligand affinity blotting, the region of rMPL36 required to bind PLG was determined by the last 70 amino acids of the protein, related to the conserved SPOR domain [36]. The aa41-305 construct bound PLG as did the intact rMPL36 (aa41-321), while the aa41-235 construct lacked binding activity (Fig 3B). Moreover, rMPL36-PLG interaction was dose-dependently inhibited by EACA, a lysine analog that binds to the PLG Kringle domains, pointing to the involvement of lysine residues in the binding of rMPL36 to PLG (Fig 3C).

**Fig 3.**
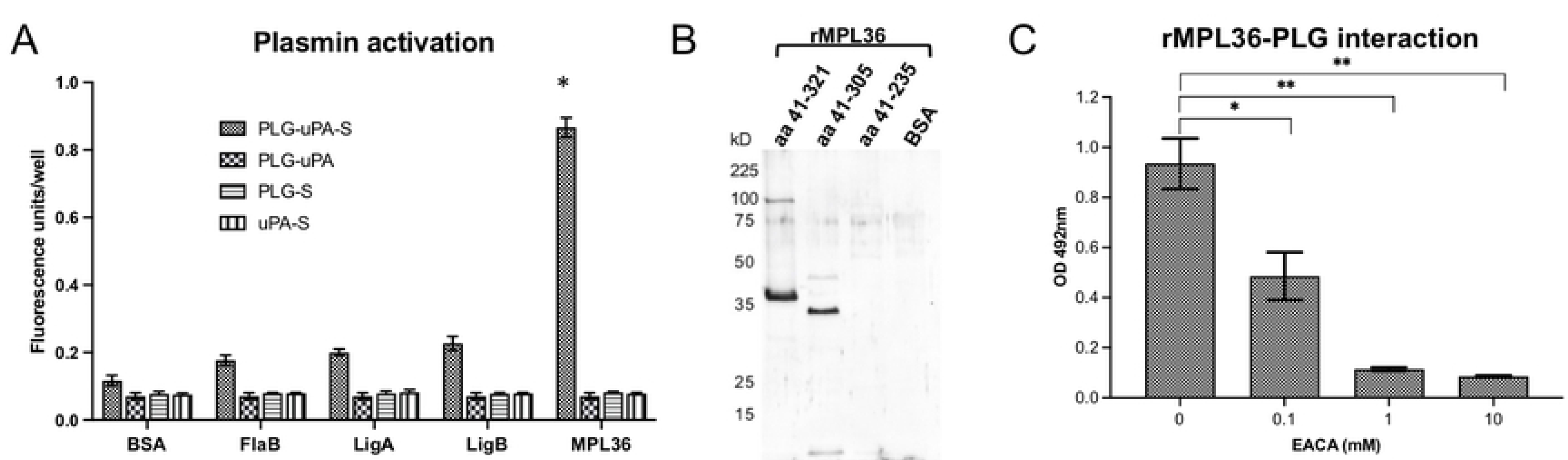
Characterization of plasminogen (PLG) binding to rMPL36. (A) MPL36-bound PLG activation to plasmin (PLA) by urokinase-type PLG activator (uPA). 96-well plates were coated with 1 µg of rMPL36, rLigA, rLigB, rFlaA2 or BSA, and incubated with PLG (1 µg). After activation with uPA (4 ng) and the chromogenic substrate (S) (0.4 mM), the reaction was measured using a fluorometer. Data represent the mean absorbance value at 405 nm ± the standard deviation of three independent experiments (\**p* < 0.001). (B) Mapping of MPL36 region interacting with PLG by ligand affinity blot. Purified full-length (aa41-321) and truncated (aa41-305 and aa41-235) MPL36 recombinant proteins were subjected to SDS-12% PAGE under reducing conditions, transferred to a nitrocellulose membrane, and incubated with 50 µg of purified human PLG. Bound PLG was detected with anti-human PLG (1:500) followed by peroxidase-conjugated anti-rabbit IgG (1:10,000). BSA was included as a negative control. (C) Assessment of lysine residues involvement in MPL36/PLG interactions. PLG (10 μg/mL) was added to MPL36-coated wells (1 μg) in the presence of epsilon-aminocaproic acid (0.1–10 mM). Bound-PLG was detected with a polyclonal anti-PLG (1:2,000), followed by peroxidase-conjugated secondary antibodies (1:10,000). Data show the mean absorbance value at 492 nm ± the standard deviation of three independent experiments (\**p* < 0.05; \*\**p* < 0.01).

The SPOR domain of MPL36 has seven lysine residues (S3A Fig). Despite differences in amino acid composition, tertiary structures of the SPOR domain from *L. interrogans* (P1), *L. fainei* (P2), and *L. biflexa* (S1) were similar (S3B Fig) and aligned with a root mean square deviation (RMSD) of 0.497. The major differences are more evident on the localization of lysine residues among different species (S3B Fig). Phylogenetic analysis revealed that the amino acid identity of MPL36 varied between 100% and 86% within pathogenic species (P1) of *Leptospira*, indicating a high degree of conservation. In contrast, the aa similarity dropped to 64% and 50% identity for pathogenic species of the P2 group and saprophyte species (S1 and S2), respectively, with MPL36 clearly being able to separate all species in different branches (S3C Fig). When we compared the amino acid composition, there were major differences between aa 248 and aa 321 among pathogenic and saprophytic species (S3D Fig), which corresponds to the region shown to be essential for PLG binding (Fig 3B).

### MPL36-bound PLG mediates fibrinogen degradation

Given the results with the rMPL36, we assessed the acquisition of human PLG in pathogenic *Leptospira* strains. The importance of MPL36 in PLG recruitment to the bacterial surface was endorsed by the reduced ability of the mutant strain to bind PLG (*p* = 0.0097), and by full restoration of PLG acquisition capacity by the complemented strain (*p* = 0.0027) (Fig 4A). Of note, binding ability of the Δ*mpl36* knockout strain to PLG was not statistically different from that displayed by the negative control strain Patoc (Fig 4A).

**Fig 4.**
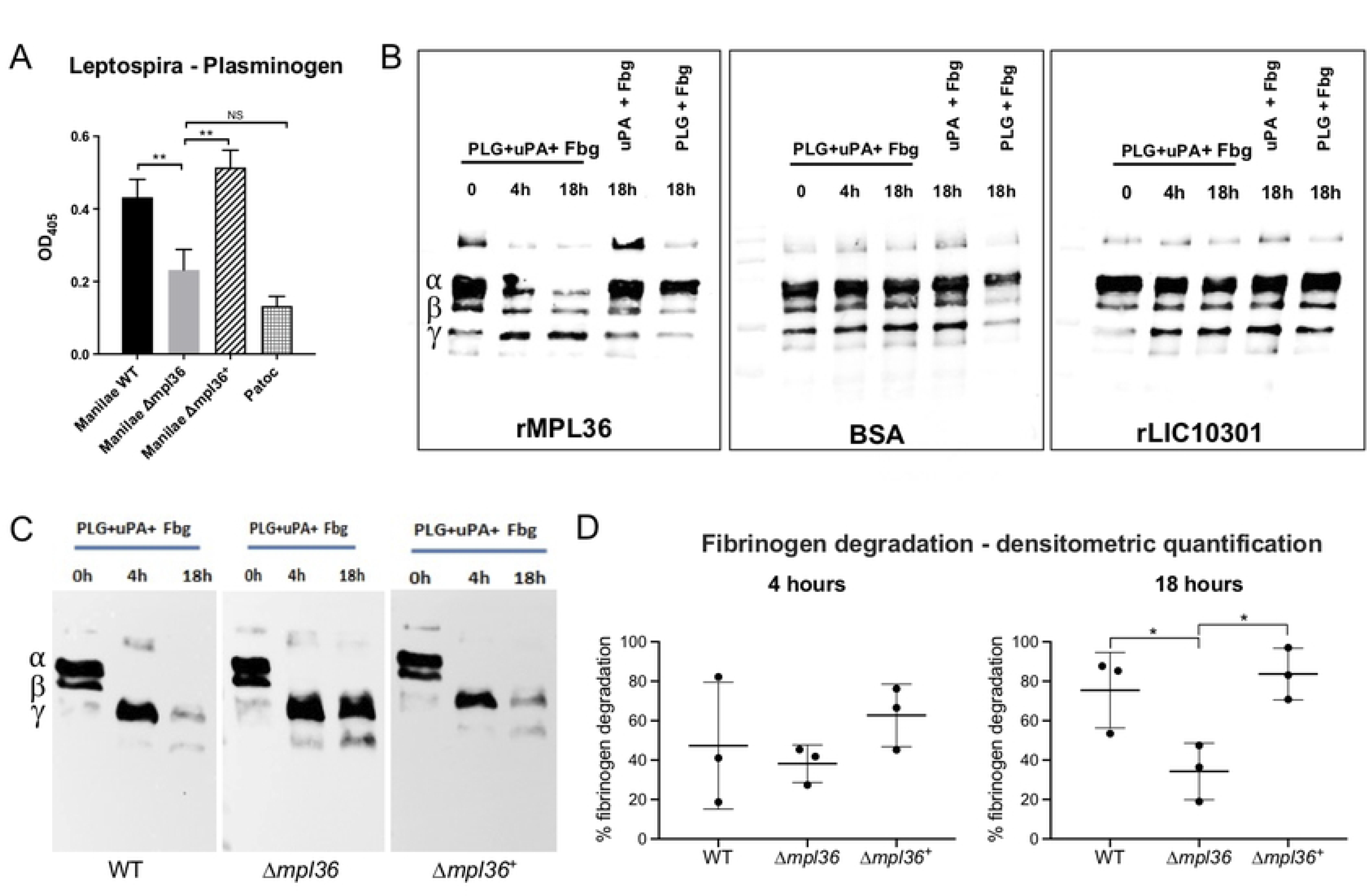
MPL36 role on the ability of *Leptospir*a cells to bind to PLG and to degrade fibrinogen (Fbg). (A) PLG binding to Manilae WT, mutant Δ*mpl36*, and Δ*mpl36*^+^ strains was assayed using a whole-cell enzyme-linked immunosorbent assay. A total of 1µg PLG was immobilized and incubated with 10^8^ bacterial cells. Bound leptospires were detected using serum from a hamster infected with an attenuated Manilae strain, and pre-immune serum was used as a control. Data represent the mean ± standard deviation of results of three independent experiments. (B) Fibrinogen degradation by immobilized rMPL36-PLA. Microtiter plate wells coated with rMPL36, rLIC10301, or BSA (10 µg/mL), were incubated with purified human PLG (20 µg/mL). After washes, fibrinogen (500 ng) and uPA (1 U) were added and incubated for up to 18 h. A western blot using anti-human fibrinogen (1:5,000) was performed. (C) Fibrinogen degradation by PLA bound to *Leptospira* strains (Manilae WT, Δ*mpl36*, Δ*mpl36*+). Leptospires (10^8^ cells) were incubated with purified human PLG (10 µg), and after washes uPA (3 U) and fibrinogen (10 µg) were added and incubated for up to 18 h. Leptospiral supernatants were collected and analyzed by western blot using anti-human fibrinogen (1:5,000). Controls excluding uPA or PLG were included. One experiment representative of three is shown. (D) Quantification of fibrinogen cleavage by *Leptospira* strains. Band intensities corresponding to α-, β-, and γ- fibrinogen chains in T0h were arbitrarily set as 100%. Quantification of fibrinogen cleavage products at T4h and T18h (relative to T0h) are shown. Data were analyzed with one-way ANOVA and Student t test (\**p* < 0.05; \*\**p* < 0.001).

We then assessed if MPL36-bound PLG, once converted to its active form PLA, could cleave ECM substrates and immune mediators of physiological importance. Recombinant protein, as well as *Leptospira* strains, were first incubated with PLG. After successive washes, preparations were incubated with fibrinogen, vitronectin, laminin, fibronectin, and C3b in the presence of the PLG activator uPA for 4 h or 18 h. rMPL36-bound PLA cleaved fibrinogen α- chain in a time-dependently manner (Fig 4B). No degradation products were detected when the plates were immobilized with the negative control proteins rLIC10301 or BSA, which do not bind PLG [30]. As expected, fibrinogen degradation was much more pronounced when we used intact bacteria. PLA bound to all three strains degraded both α and β chains of fibrinogen.

However, almost complete degradation was only achieved in the presence of the WT and the complemented strains after 18 h incubation period (Fig 4C).

Fibrinogen degradation in the presence of both the WT and complemented strains at 18 h was significantly more efficient (*p* = 0.0447 and *p* = 0.0119, respectively) compared to the one observed in the presence of the mutant strain (Fig 4D). As ECM and complement proteins are among PLA targets, we also assessed the ability of MPL36-bound PLA to degrade vitronectin (S4A Fig), laminin (S4B Fig), and complement C3b (S4C Fig). No differences regarding the degradation patterns were observed for the WT, Δ*mpl36* and Δ*mpl36+* strains (S4A-C Fig), thus indicating that MPL36-bound PLA specifically targets fibrinogen. Complete degradation of vitronectin probably results from the action of bacterial secreted proteases since the cleavage profiles are quite similar for all three strains (S4A Fig). Taken together, those results indicate that MPL36 is a major PLG binding protein of pathogenic leptospires and contributes to the generation of a proteolytic platform on the leptospiral surface, which could increase the invasiveness potential of this bacterium.

### Native MPL36 promotes *Leptospira* adhesion to epithelial host cells and translocation

To further confirm the adhesive and invasiveness properties of MPL36 on epithelial cells, the interaction of Manilae WT, Δ*mpl36*, and Δ*mpl36^+^* strains with cultured MDCK cells was investigated. The saprophyte Patoc strain was included as a negative control. An average of 6 WT leptospires adhered to each MDCK cell while approximately 0.2 and 0.5 of mutant and saprophytic leptospires, respectively, were found associated with each MDCK cell (Fig 5A). The binding capacity of the mutant strain was significantly reduced compared to the ones displayed by the WT (*p* = 0.001) and the complemented strain (*p* = 0.0107), but the phenotype of adherence was not completely restored on the complemented strain (average of 4 leptospires/MDCK cell) when compared to the WT (*p* = 0.0452, Fig 5A).

**Fig 5.**
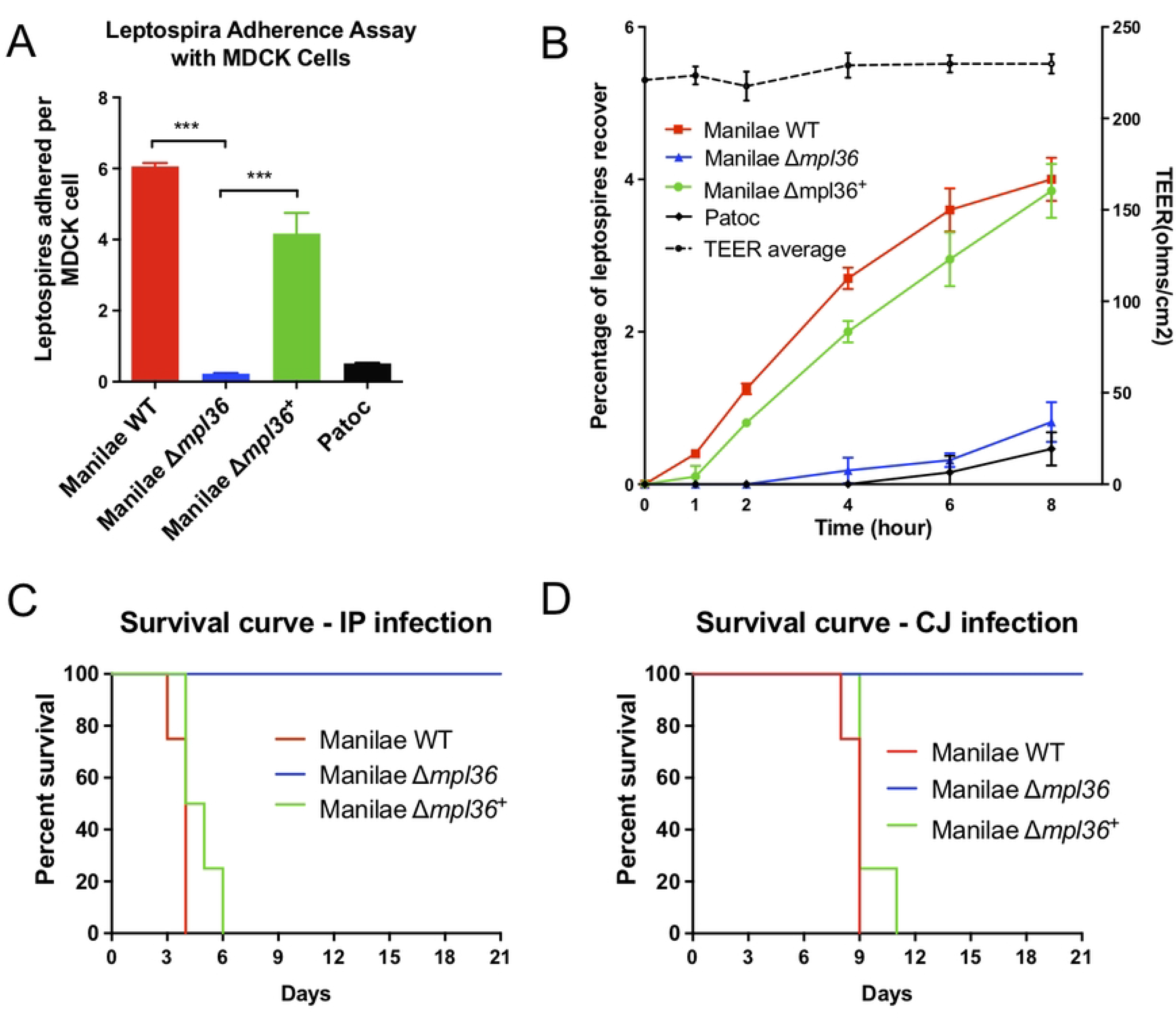
Role of MPL36 on the pathogenesis of *Leptospira interrogans*. (A) Ability of leptospires strains to adhere to MDCK epithelial cells. Manilae WT, Δ*mpl36*, and Δ*mpl36*^+^ strains were incubated with immobilized MDCK cells for one hour at 37°C and adhesion assessed by immunofluorescence. Data represent the mean ± standard deviation of results from three independent experiments. (B) Ability of *Leptospira* strains to translocate through polarized monolayers of MDCK cells. The Patoc strain was used as negative control. Strains were inoculated (100 MOI) in the upper chamber of a Millicell culture plate containing MDCK cell monolayer. Percent recovery of leptospires was determined by counting bacteria in the lower chamber between 0 and 8 hours after inoculation. Left Y axis shows TEER measurements. Data represents the mean ± standard deviation of results from three independent experiments. (C, D) Survival curve of hamsters infected with *Leptospira* strains. Animals were infected either by intraperitoneal (IP) or conjunctival (CJ) route with 10^8^ leptopires. Data represent results of one of two independent experiments.

We also investigated the ability of these *Leptospira* strains to translocate through monolayers of MDCK cells. Within the first two hours after infection, only the WT and complemented strains were recovered from the lower chamber of the Millicell culture plates, indicating their ability to successfully cross the MDCK cell monolayers (Fig 5B). The mutant strain (Δ*mpl36)* was recovered starting at 4 h post-infection. However, over the 8-hour course of the experiment, there is a statistical difference in the number of mutant cells that were recovered compared to the WT (*p* = 0.0284) and complemented strains (*p* = 0.0456). There were no statistical differences between the numbers of WT and complemented strains recovered (*p* = 0.0625), indicating that the complementation restored the ability of the mutant to translocate. As expected, the Patoc strain was unable to successfully translocate, as previously observed [40] (Fig 5B). Consistent TEER measurements indicated that disruption of cell monolayers’ TJ did not occur during the translocation process (Fig 5B). Taken together, those experiments indicate that MPL36 abrogation partially impairs the ability of leptospires to adhere to and translocate through polarized monolayers of MDCK cells, which in turn would affect the ability of the pathogen to efficiently disseminate and cause disease.

### MPL36 is essential for virulence of pathogenic *Leptospira* during acute infection

To determine the role of MPL36 during the infection process, the WT, Δ*mpl36* and Δ*mpl36*^+^ strains were tested for virulence in the hamster model of leptospirosis infection. Hamsters challenged by IP route with 10^8^ leptospires of the *Δmpl36* strain survived up to 21 days without any symptoms of infection, whereas hamsters challenged with the same number of bacteria of the WT strain died between 3-5 days post-infection (Fig 5C and S2 Table). The mutant phenotype complemented by reintroduction of *mpl36* yielded 100% lethality at dose 10^8^ in 4-6 days after IP challenge (Fig 5C and S2 Table).

When using the CJ route of infection, hamsters challenged with 10^8^ leptospires with the Δ*mpl36* mutant survived without disease manifestation, whereas all animals challenged with the WT strain died between days 8-9 post-infection (Fig 5D and S2 Table). Animals infected with the complemented strain all died between 9- and 11-days post-infection (Fig 5D and S2 Table). Leptospires were not detected in the kidney of animals infected with Δ*mpl36* by both routes of infection at day 21 post-infection, when animals were euthanized (S2 Table). Those results indicate that MPL36 is an essential protein for the pathogenesis of *Leptospira* (Fig 6).

**Fig 6.**
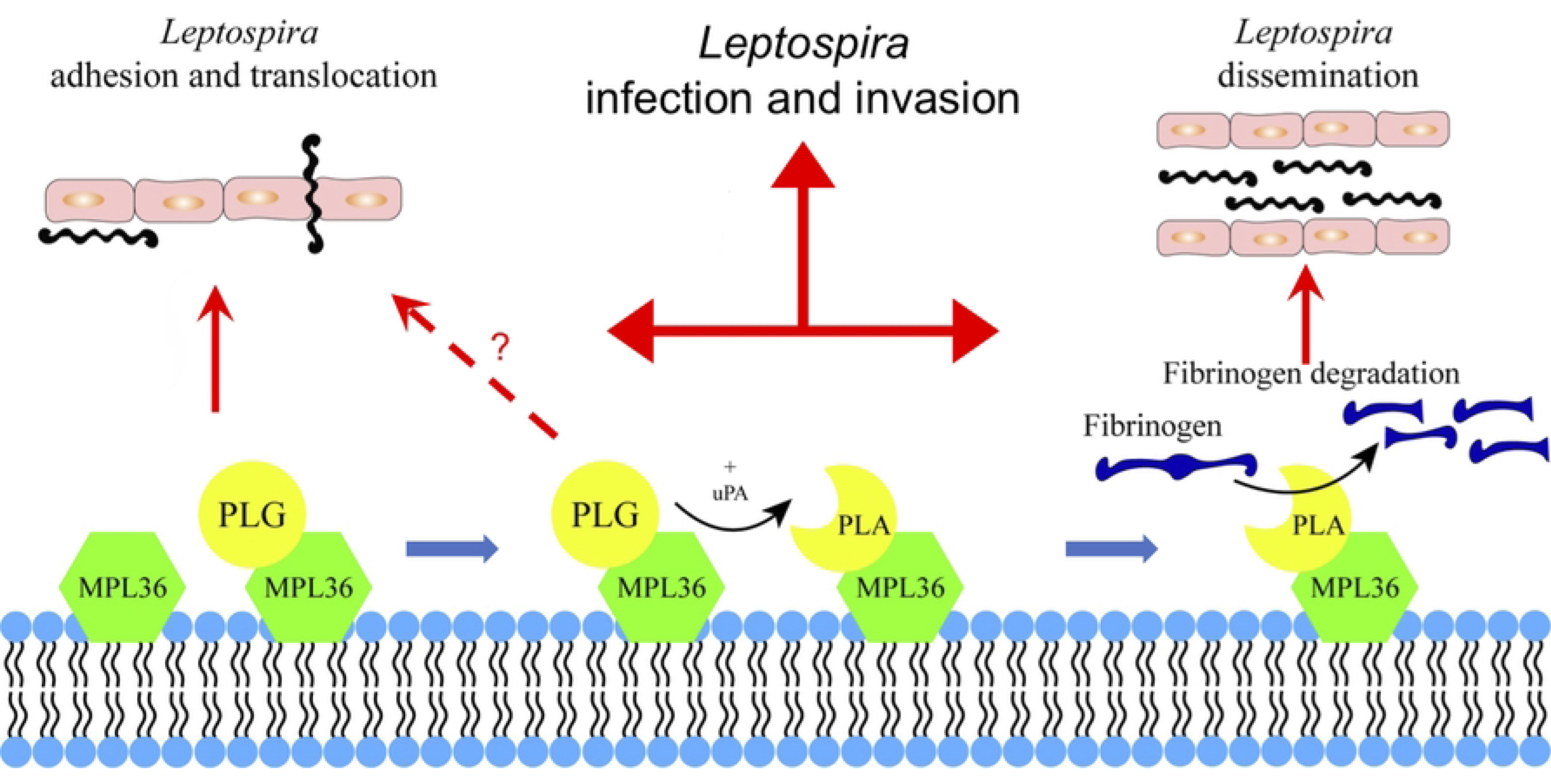
**Schematic summary view of MPL36 role on leptospiral pathogenesis**. Hypothetical model showing how acquisition of PLG by MPL36 contributes to *Leptospira* adhesion and translocation through cell monolayers, and to the proteolytic potential of this spirochete.

## Discussion

Our results showed that MPL36 is an outer membrane protein acting as a major PLG binding in pathogenic *Leptospira*, with the ability to degrade fibrinogen, being essential for leptospiral virulence. The mechanisms behind the pathogenesis of leptospiral infection continues to be a major knowledge gap preventing the advance on diagnostic and prevention. Despite the recent major breakthroughs on the research of this important neglected zoonotic disease [21], more work is needed to better understand how leptospires can rapidly disseminate, evade host-immune responses, and cause life-threatening disease worldwide. A previous study has shown that only virulent strains of *Leptospira* spp. were capable of acquiring PLG from human plasma, strongly suggesting the involvement of this process in leptospiral virulence [27]. Through capturing PLG on their surface, followed by activation to PLA, these spirochetes augment their proteolytic capacity, and consequently, potentiate dissemination through tissue barriers. It has also been suggested that direct binding of fibrinogen to the leptospiral surface could lead to an increased consumption of those molecules, causing a reduction in fibrin clot formation [41].

*Leptospira* proteome contains multiple proteins with the ability to bind different ECM and PLG [4, 32]. However, those studies were conducted using recombinant proteins and none of those targets have been confirmed to have a direct role in virulence.

The leptospiral MPL36 belongs to a family of proteins containing a Rare lipoprotein A (RlpA) domain [42] and a C-terminal SPOR (sporulation related repeat) domain [36]. Similar to its homologous in *E. coli* and *P. aeruginosa* [38, 43], MPL36 has been characterized as a putative surface exposed protein in *Leptospira* [28, 44]. The gene *mpl36* has been shown to be upregulated in a dialysis membrane chamber (DMC) system implanted in the peritoneal cavities of rats [45], representing a mammalian host-adapted state, and it has been identified as one of the genes regulated by *Leptospir*a virulence regulator (*lvrAB*) [46], a signaling system that controls virulence in *Leptospira*. More recently, MPL36 has been identified as one of the targets associated as correlates of cross-protection on an attenuated-vaccine model for leptospirosis [47]. Furthermore, we demonstrated that human patients with leptospirosis produced antibodies that recognize MPL36 [44], and that IgM antibodies against MPL36 are detected during the early stages of the disease, thus indicating that MPL36 is expressed during host infection, has an important role on pathogenesis, and might serve as a target for serodiagnosis and vaccine development.

MPL36 was first characterized as a PLG-binding molecule in a study aiming to identify *Leptospira* PLG receptors [28]. In this study, we explored further the interaction of MPL36 with PLG by evaluating the functional consequences of PLG acquisition by leptospires through this surface receptor as well as leptospiral virulence in the absence of *mpl36*. Our results with both recombinant protein and mutant *Leptospira* strains not only confirmed the ability of MPL36 to bind to host epithelial cells, but also demonstrated that a mutant unable to express MPL36 (Δ*mpl36*) had an impaired binding capacity to PLG, similar to the phenotype observed for the saprophyte *L. biflexa*. The process of PLG binding to bacterial receptors on the cell surface has been shown for several pathogens to be mediated by the PLG Kringle domains containing lysine-binding sites [1]. Our experiments mapped the PLG-interacting region of MPL36 to the SPOR domain on the protein C-terminus, and binding was shown to occur through lysine residues within this domain.

SPOR domains are about 70 amino acids long consisting of four antiparallel β-sheet flanked on one side by two α-helices. The domain primary structure is not highly conserved, but it is present in several bacteria as a peptidoglycan (PG) binding domain and its function is mainly related to remodeling the PG sacculus during cell division [36, 37]. Most bacterial SPOR domain proteins are present in the periplasm, although both *P. aeruginosa* and *E. coli* RlpA proteins are outer membranes [35, 38]. *P. aeruginosa* outer membrane RlpA was described as a lytic transglycosylase [38], with mutants having striking morphological defects. However, none of the *E. coli* SPOR proteins have been reported to have an enzymatic activity [48] or being related to growth or cell viability [35]. MPL36 binding to PG and participation in cell division were not directly assessed in this study. However, the Δ*mpl36* strain had no morphological or growth defects, strongly suggesting that MPL36 is not involved in these processes in *Leptospira*.

Although found in all *Leptospira* species, with 50% amino acid identity in the saprophyte *L. biflexa*, our *in-silico* analysis disclosed differences in primary structure composition, mainly clustered on the SPOR domain of the protein, suggesting a potential different function of this protein in non-pathogenic species.

MPL36-mediated PLA formation degrades fibrinogen on pathogenic leptospires. A partial yet significant degradation of fibrinogen was observed in the presence of rMPL36, suggesting the existence of additional PLG-interacting proteins on the leptospiral surface able to degrade fibrinogen, as previously reported [4, 26, 32, 49]. This assumption is further supported by the fact that *Leptospira* binding to PLG, although significantly reduced, was not completely abolished when we used the Δ*mpl36* mutant strain. It is also important to highlight that fibrinogen degradation by *Leptospira* results from two distinct mechanisms: PLA-binding activity and proteolytic action of extracellular metalloproteases secreted by virulent strains, which target ECM and plasma proteins from the host [25]. The combined strategies ensure an efficient degradation of host fibrinogen by *Leptospira*. However, after 18 h of incubation almost complete hydrolysis of this coagulation cascade molecule was observed only when both WT or Δ*mpl36^+^*strains were incubated with PLG, uPA and fibrinogen. Curiously, our experiments showed that MPL36-bound PLA had no apparent effect on vitronectin, laminin or complement C3b. Taken together, the MPL36-bound PLA seems to have an essential and targeted role on fibrinogen degradation, and more studies are needed to confirm and better understand this peculiar activity.

MPL36 mediates binding of leptospires to MDCK cells, a crucial step for subsequent translocation through cell monolayers. Studies have shown that in *Streptococcus* sp. the adherence process for colonization and translocation were dependent of PLG recruitment in the surface, with PLG acting as a linker molecule independently of the plasmin activity [50, 51]. rMPL36 did not bind purified fibronectin or laminin, indicating the possibility that MPL36- bound PLG could anchor leptospires to the cell surface, serving as a bridging molecule. There is no evidence of ECM degradation related to MPL36, thus the inability of Δ*mpl36* strain to translocate could be strongly related to the low capability to adhere to host cells. Nevertheless, lack of expression of MPL36 in pathogenic *Leptospira* lead to a complete attenuation phenotype in hamsters, challenged by both IP and CJ routes, with the latter mimicking a more natural route of infection that involves adhesion and translocation of host cells. Bacterial PLG receptors are described as multifunctional surface proteins during infection process [1], and it has been shown that lysine residues must be precisely positioned during presentation to substrates and catalysts in order to activate the plasminogen system [52], thus influencing the pathogenic process. For that reason, further studies are necessary to investigate other potential roles that MPL36 might have on the leptospiral pathogenesis process, leading to the observed attenuated phenotype.

In this study, we characterized the MPL36 protein as a surface exposed virulence factor, with the ability to bind to PLG and degrade fibrinogen, playing a role on adhesion and subsequent translocation of the leptospiral spirochete on host cells (Fig 6). Genetic manipulation tools for targeted mutagenesis in *Leptospira* are still limited and restricted to a few research laboratories, and the identification and characterization of leptospiral proteins’ interaction with host components have been mostly based on recombinant proteins. Our results are based on *in vitro* and *in vivo* analysis using knockout and complemented mutants, fulfilling molecular Koch’s postulates, providing new insights regarding bacterial proteins that explore PLG for their pathogenesis. There is a major interest in the identification of antigenically conserved surface-exposed proteins with the capacity to serve as broader vaccine candidate targets. In addition, the characterization of leptospiral components contributing to pathogenesis would aid in the development of improved diagnostic strategies. Our results demonstrate that MPL36 is a highly conserved protein among pathogenic species, expressed and able to elicit immune response during the infection process, satisfying all the requirements to be further explored on research designed to better understand leptospiral pathogenesis and advance diagnostic and prevention of this important zoonotic disease.

## Material and methods

### Bacterial strains, cells, and culture conditions

The pathogen *Leptospira interrogans* serovar Manilae strain L495 (Manilae WT), the Manilae *mpl36* mutant (Δ*mpl36*), the Manilae *mpl36* mutant complemented strain (Δ*mpl36^+^*) and the saprophyte *L. biflexa* serovar Patoc strain Patoc 1 (Patoc) were grown in Ellinghausen-McCullough-Johnson-Harris (EMJH) liquid medium [53] with agitation or on 1% agar plates at 30°C. *E. coli* cells were grown in Luria-Bertani (LB) medium at 37°C with agitation. When appropriate, spectinomycin and/or kanamycin were added to the culture medium at a final concentration of 50 μg/mL. Madin–Darby canine kidney (MDCK) cells (ATCC CCL34TM) were cultured in minimum essential medium (MEM) containing 10% fetal bovine serum (GIBCO Laboratories). Cells were grown at 37°C in a humidified atmosphere with 5% CO2.

For growth curves of WT, mutant and complemented strains, bacteria were initially enumerated under dark-field microscopy by using Petroff-Hausser counting chambers (Fisher Scientific) and diluted to a starting bacterial concentration of 10^4^ mL at 30°C. Growth was monitored daily by counting. Three independent experiments were performed. For the motility assay, 5 μL of 10^5^ leptospires were inoculated on 0.5% agarose plates and incubated for 10 days at 30°C, as previously described [40]

### Proteinase K treatment of intact bacteria

MPL36 cell surface localization was first assessed by proteinase K treatment as previously described [39]. Briefly, Manilae WT cells were grown to a density of 5×10^8^ cells/mL and harvested by low-speed centrifugation at 2,000 × *g* for 10 min at room temperature. The pellet was gently washed with phosphate buffered saline (PBS) containing 5 mM MgCl2, and collected by centrifugation at 2,000 × *g* for 10 min. After resuspension in PBS-5 mM MgCl2, proteinase K (Sigma-Aldrich) diluted in proteolysis buffer (10 mM Tris-HCl pH8.0, 5 mM CaCl2) was added to the washed leptospires in a final concentration of 25 to 100 µg/mL. Proteolysis buffer without proteinase K was added to the negative control. The reaction was quenched by the addition of phenylmethylsulfonyl fluoride (PMSF). Leptospires were subsequently collected by centrifugation and washed twice with PBS-5 mM MgCl2, and the cells were resuspended in sample buffer for SDS-PAGE. Immunoblot analysis was performed using rabbit antibodies against proteins MPL36, LigA and GroEL at a dilution of 1:1,000. Bound antibodies were detected using horseradish peroxidase (HRP)-conjugated anti-rabbit IgG (GE Lifesciences) at a dilution of 1:100,000. Positive signals were detected by SuperSignal™ West Pico Kit (Pierce), according to the manufacturer’s instructions, and blots were analyzed using ChemiDoc™ Imager (Bio-Rad).

### Immunogold labeling

Bacterial cultures (1 x 10^8^ cells/mL) were centrifuged (6,500 x *g* for 20 min), washed with PBS three times and fixed with 4% formaldehyde for 30 min. After fixation, preparations were washed three times with PBS and blocked with 0.2% bovine serum albumin (BSA) in PBS (PBS-BSA) for 30 min. Preparations were then incubated overnight with rabbit anti-MPL36 antiserum (1:10 dilution in PBS) at 4°C. Subsequently, preparations were washed with PBS-BSA, and incubated with goat anti-rabbit antibody labeled with 10 nm colloidal gold particles (Sigma-Aldrich) diluted 1:10 in PBS, for 4h at room temperature. After further washings, preparations were mixed (1:1) with 2% uranyl acetate (UA) in water and placed onto Formvar-coated nickel grids for 2 min. After completely dried with filter paper, preparations were then analyzed under TEM (LEO 906E – Zeiss, Germany) at 80 kV.

### Phase partitioning of *Leptospira* membrane proteins using Triton X-114

Phase separation of the integral membrane proteins of *Leptospira* to localize the protein MPL36 was performed using Triton X-114 solution as described elsewhere [54]. Briefly, a 50 mL mid-log-phase culture of *L. interrogans* serovar Copenhageni Fiocruz L1-130 (5×10^9^ cells) was washed in PBS containing 5 mM MgCl2. The membrane proteins were extracted at 4°C with 1% Triton X-114, 150 mM NaCl, 10 mM Tris (pH 8.0) and 1 mM EDTA. The insoluble debris was removed by centrifugation at 12,000 × *g* for 15 min and then 20 mM CaCl2 was added to the supernatant. Phase separation was performed by warming the supernatant at 37°C and subjecting it to centrifugation for 10 min at 1,000 × *g*. The detergent and aqueous phases were separated and precipitated with 10 volumes of chilled acetone. The aqueous and detergent phases were then resolved onto 12% SDS-polyacrylamide gel before transferring to nitrocellulose membrane for immunoblot analysis using polyclonal rabbit sera anti-MPL36, anti-LigA, and anti-GroEL (1:1,000).

### Cloning, expression, and purification of MPL36 fragments

The full recombinant MPL36 protein (rMPL36/aa 41-321) was commercially produced by GenScript^®^ Biotech with His-tag in an *E. coli* expression system. Recombinant MPL36 protein devoid of the last 16 residues (rMPL36/aa 41-305) and devoid of the last 86 residues (MPL36/aa 41-235) were produced following protocol previously described [39]. DNA fragments were amplified by PCR from *L. interrogans* serovar Copenhageni genomic DNA (strain Fiocruz L1-130) with set of primers aa41-305 and aa41-235, using restrictions enzyme sites for BamHI and NcoI, respectively (S1 Table). The amplified products were cloned into the pGEM-T Easy vector (Promega) and subcloned into the pAE expression vector [55]. This vector allows the expression of recombinant proteins with a minimal His6 tag at the N-terminus. The constructs were transformed into *E. coli* BL21 DE3. Expression and purification of the recombinant proteins were performed essentially as previously described [39].

### Binding assay using fluorescent latex beads

To assess the binding ability of rMPL36 to ECM proteins and PLG, l µg of fibronectin, laminin, PLG (Sigma-Aldrich) and BSA (negative control) was coated on 96-well plates overnight at 4°C. Plates were blocked for 1 h at 37°C with 5% non-fat dried milk in 2% BSA. Recombinant proteins MPL36, FlaA2 (negative control), LigA and LigB (positive controls) were diluted in PBS to a final concentration of 0.2 nmol. A 4 µL sample of the stock suspension of latex beads (containing about 3×10^10^ mL^-1^ of 0.3 µm diameter beads, Sigma-Aldrich) was added to each protein and incubated for 2 h at 37°C. A sample of 100 µL of this suspension was then added in triplicate to the wells of the previously coated plates and incubated for 2 h at 37°C. Plates were washed four times with PBS-0.05% (vol/vol) Tween 20 (PBST) and the fluorescent emission (excitation at 486-580 nm and emission at 568-590 nm) was measured using Synergy HT (BioTek, Agilent). Uncoated beads and beads coated with BSA were used as controls. For statistical analyses, the experiment was repeated three times and the attachment of recombinant proteins to the host components was compared to its binding to BSA.

To assess the binding ability of rMPL36 to host epithelial cells, Madin-Darby Canine Kidney (MDCK) cells were seeded in a 24-well tissue culture plate at a density of 2 x 10^5^ cells per well and incubated at 37°C with 5% CO2 for 48 h until the formation of a near-confluent MDCK cell monolayer. Samples of 100 µL of the latex bead suspension coated with rMPL36, rFlaA2, rLigA and rLigB, as described above, were added in triplicate to the pre-incubated MDCK cells. After 5 h incubation at 37°C, cells were washed three times with PBS. The fluorescence emission was measured directly in the 24-well plate using Synergy HT (BioTek, Agilent). The experiment was repeated twice.

### Conversion of PLG into active PLA

To evaluate if PLG bound to rMPL36 is converted into active PLA 96-well plates were coated with 0.2 µM of recombinant proteins or BSA (negative control) at 4°C overnight. The plates were washed once with PBST and blocked 2 h at 37°C with 10% non-fat dried milk in PBS. The blocking solution was discarded, and the plate washed four times with PBST. Then, 1 µg of human PLG (Sigma-Aldrich) was added to each well and incubated for 2 h at 37°C. Wells were washed four times with PBST, and then 4 ng of human urokinase-type PLG activator (uPA, Sigma-Aldrich) was added per well. After that, 100 µl of the PLA specific substrate D-valyl- leucyl-lysine-p-nitroanilide dihydrochloride (Sigma-Aldrich) was added at a final concentration of 0.4 mM in PBS. Plates were incubated overnight, and substrate degradation was measured at 405 nm using Synergy HT (BioTek, Agilent). The experiment was repeated three times for reproducibility.

### Role of lysins in rMPL36-PLG interaction

To assess the role of lysins in rMPL36-PLG interactions 96-well plates were coated with rMPL36 (10 µg/mL). After blocking with 3% BSA, PLG (10 µg/mL) and ε -aminocaproic acid (0 – 10 mM) were added to the coated wells. Bound PLG was detected with a rabbit polyclonal antibody (Sigma-Aldrich) at a 1:2,000 dilution followed by peroxidase-conjugated anti-rabbit IgG (Sigma-Aldrich) at a 1:10,000 dilution. Student’s two-tailed *t* test was used for statistical analyses.

### Interaction of rMPL36 with PLG by ligand affinity blotting

Purified recombinant proteins were subjected to 12% SDS–PAGE and transferred to a nitrocellulose membrane. After blocking with 5% BSA the membrane was incubated for 1 h with 50 µg of purified PLG (Sigma-Aldrich) diluted in PBS. After five washes with PBST, the membrane was incubated with polyclonal rabbit antibodies recognizing human PLG (Sigma-Aldrich, 1:500), followed by peroxidase-conjugated secondary antibodies (1:10,000). Positive signals were detected by SuperSignal™ West Pico Kit (Pierce). BSA was used as a negative control.

### Detection of antibodies to rMPL36 using human sera of individuals with laboratory-confirmed leptospirosis

We performed an ELISA assay using rMPL36 against acute and convalescent sera from 29 individuals with confirmed severe leptospirosis enrolled in our surveillance study in Salvador, Brazil [14]. Control human sera were obtained from healthy US human donors. Microtiter plates (Corning) were coated with 80 ng of rMPL36 and incubated overnight at 4°C. The plates were washed three times with PBST and incubated with 5% milk blocking solution for 2 h at 37°C. After four washes with PBST, wells were incubated in duplicate with human immune sera, diluted 100-fold in 2% BSA, for 1 h at 37°C. Secondary anti-IgM human HRP conjugated antibody (Sigma-Aldrich) or anti-IgG human HRP (Jackson ImmunoResearch) was used at a dilution of 25,000 (2% BSA) and incubated for 1 h at 37°C. Tetramethylbenzidine (TMB, SureBlue Reserve) was used for detection and the reaction was stopped by adding 100 μL of 2 N H2SO4. Absorbance (450 nm) was recorded using Synergy HT (BioTek, Agilent). A threshold was calculated based on 2.5 SD of the average OD from the healthy individuals for each isotype.

### Insertion mutagenesis and complementation

Random mutagenesis in *L. interrogans* serovar Manilae strain L495 using mariner-based transposon *Himar1* has been previously described [33]. The insertion site was identified by semi-random PCR followed by DNA sequencing. The insertion within *mpl36* was confirmed by PCR using primers flanking the insertion site. For complementation, *mpl36* and its native promoter were PCR amplified from Manilae WT using primers mpl36F and mpl36R, designed with KpnI restriction sites (S1 Table). The amplicon was then digested by KpnI and ligated into plasmid pAL614, which carries a modified *Himar1* transposon for conjugation [56] containing a spectinomycin resistance cassette. The resulting plasmid (pAL614-*mpl36*) was used to chemically transform *E. coli* S17 cells, which were subsequently used to transform serovar Manilae Δ*mpl36* strain via conjugation. Transposon insertion occurred at nucleotide 292,919 in the gap of two open reading frames. MPL36 expression by the mutant (Manilae Δ*mpl36*) and complemented (Manilae Δ*mpl36^+^*) strains was verified by Western-blot analysis using anti- MPL36, as described above.

### Binding of *Leptospira* strains to PLG

Adhesion of *Leptospira* strains (Manilae WT, Δ*mpl36*, Δ*mpl36^+^*, and Patoc) to human PLG (Sigma-Aldrich) was assessed by ELISA as previously described [23]. A 96-well plate was coated with PLG or BSA (negative control) (10 µg/mL) overnight at 4°C. Plates were washed four times and then blocked with 5% non-fat dried milk in 2% BSA for 1 h at 37°C. After washing with PBST, 1×10^8^ leptospires were added to each well and incubated for 90 min at 37°C. After two washes with PBST to eliminate non-adherent cells, adherent leptospires were fixed with cold-methanol for 10 min at -20°C, and detected with a hamster polyclonal serum of animals infected with 10^7^ *L. interrogans* serovar Manilae *fcpA^-^*[40] (1:1,000) followed by a peroxidase-conjugated anti-hamster IgG (1:50,000). Pre-immune serum was used as a negative control, and final values were calculated from three technical results after subtracting the pre-immune sera data. Absorbance was measured at 405 nm using Synergy HT (BioTek, Agilent).

### ECM and plasma protein degradation by PLA bound to *Leptospira* strains

Mid-log phase leptospires (Manilae WT, Δ*mpl36*, Δ*mpl36^+^*) were harvested by centrifugation at 9,000 x *g* for 5 min and 1 x 10^8^ bacteria were incubated with purified human PLG (10 µg) diluted in PBS for 1 h at 37°C. Leptospires were washed three times with PBS and then incubated with 3 U of uPA (Sigma-Aldrich) and 10 µg of human PLG-depleted fibrinogen (Calbiochem), or 2.5 µg of human vitronectin (Sigma-Aldrich), or 5 µg of laminin (derived from mouse Engelbreth-Holm-Swarm sarcoma, Sigma-Aldrich), or 2.5 µg of human plasma fibronectin (Sigma-Aldrich), or 1.5 µg of human complement C3b (Complement Technology) at 37°C for 0, 4 and 18 h. For the control reactions, either PLG or uPA was omitted. After incubation, leptospiral supernatants were collected by centrifugation and subjected to 12% SDS-PAGE. The cleavage fragments were analyzed by western blot with either mouse anti-human fibrinogen polyclonal antibody (Sigma-Aldrich) at a 1:5,000 dilution, rabbit anti-human vitronectin polyclonal antibodies (Complement Technology) at a 1:5,000 dilution, rabbit anti-human laminin polyclonal antibodies (Sigma-Aldrich) at a 1:5,000 dilution, rabbit anti-human fibronectin polyclonal antibodies (Sigma-Aldrich) at a 1:5,000 dilution, or goat anti-human C3 polyclonal antibodies (Complement Technology) at a 1:5,000 dilution, followed by peroxidase-conjugated anti-mouse (1:5,000), rabbit or goat (1:10,000) IgG (Sigma-Aldrich). Positive signals were detected by SuperSignal™ West Pico Kit (Pierce). Images of the three experiments performed were recorded and band intensities were quantified using the Alliance LD2 system (Uvitec, Cambridge, UK). Band intensities at 0 h, corresponding to α-, β-, and γ- fibrinogen chains, were arbitrarily set as 100%.

Degradation of fibrinogen by rMPL36 was also assessed. rMPL36, and negative controls rLIC10301 [30] and BSA (10 μg/mL) were immobilized on microtiter plate wells and blocked with 3% BSA diluted in PBS. PLG (20 μg/mL) was added, and plates were incubated for 1 h at 37° C. Wells were then washed six times with PBST and added with human fibrinogen (500 ng/well, PLG depleted; Calbiochem) and uPA (1 U/well). Reaction mixtures were incubated at 37° C for the indicated time points and were then separated by 12% SDS-PAGE. The degradation products of fibrinogen were detected by Western blot using rabbit anti-human fibrinogen polyclonal antibodies (Sigma-Aldrich) at a 1:5,000 dilution as described above.

### Adherence to and translocation of leptospires through MDCK cells

Adherence of leptospires to MDCK cells was assessed as previously described [23].

MDCK cells (1 x 10^5^) were added to round coverslips in 24-well plates and incubated until the formation of a near-confluent MDCK cell monolayer. *Leptospira* cells were incubated with MDCK at a multiplicity of infection (MOI) of 100 for 1 h at 37°C in 5% CO2. Following incubation, the suspensions were removed, and the monolayers were washed six times with warm PBS. The cells were fixed with pre-cold (-20°C) methanol for 10 min at 4°C, followed by three washes. Blocking buffer (10% fetal bovine sera in PBS) was then added to each well and incubation proceeded for 1 h at 37°C. To detect the adherent cells, a rabbit polyclonal antiserum against the protein LipL32 of *Leptospira* was used as primary antibody at a concentration of 1:200, followed by goat anti-rabbit antibodies conjugated with Alexa488 (Jackson ImmunoResearch) both at a concentration of 1:200 diluted in blocking buffer. Incubations were performed at 37°C for 1 h followed by three washes with PBS. After the final wash, the coverslips were removed from the 24-well plates, stained with DAPI, and mounted on a glass slide. Fluorescent-labeled leptospires associated to 100 MDCK cells on each of three poly-D-Lysine treated glass coverslips were enumerated. These three coverslips served as technical replicates for each strain tested during each trial.

Bacterial translocation across epithelial cells was assessed, as previously described [40].

MDCK cells (2 x 10^5^ cells) were cultured on polycarbonate Millicell culture plate inserts (12- mm diameter, 3-µm pore size; Merck Millipore) at 37°C under 5% CO2 atmosphere.

Transepithelial electrical resistance (TEER) of the filter-grown monolayers was measured using a Millicell-ERS device (Merck Millipore) as an index for integrity of the tight junctions (TJ).

Polarized monolayers exhibiting TEER values of 150-300 Ωcm^2^ were used as an *in vitro* model of the epithelial barrier. The monolayers were infected with leptospires at a MOI of 100. The ability of *Leptospira* strains to translocate across the epithelial barrier was assessed by quantifying bacteria in the culture medium recovered from the lower chambers at 1, 2, 4, 6, and 8 h after infection.

### Evaluation of virulence in hamster model of infection

The virulence of Manilae WT and the mutant strains (Δ*mpl36* and Δ*mpl36^+^*) was assessed in 3-week male Golden Syrian hamsters, as described previously [40]. Groups of 3 hamsters were infected via intraperitoneal (IP) and ocular conjunctiva (CJ) routes with 10^8^ leptospires.

When appropriated, a LD50 experiment was performed using 10^8^, 10^6^ and 10^4^ leptospires by IP route or 10^8^ and 10^6^ by CJ route. Experiments were repeated at least once for reproducibility. Animals were monitored twice daily for signs of disease and death, up to 21-days post-infection. Surviving animals at the end of the experiment or moribund animals showing difficulty in moving, breathing, and/or signs of bleeding or seizure were immediately sacrificed by inhalation of CO2. Blood, liver, and kidneys were collected from all animals after euthanasia. The protocol of animal experimentation was prepared and approved according to the guidelines of Institutional Committee for the Use of Experimental Animals, Yale University (protocol # 2023-11424).

Kidneys were harvested and DNA was extracted for quantitative real-time PCR (qPCR) targeting *lipL32*, as previously described [40].

### Bioinformatic and phylogenetic analysis

Tertiary structure prediction of the SPOR domains from RlpA of *L. interrogans*, *L. fainei* and *L. biflexa* tertiary structure prediction was determined by AlphaFold [57], and PyMOL software (version 2.5.4) was used for analysis. Sequences of MPL36 SPOR domain of *L. interrogans* and from 69 different species of *Leptospira* spp., representing pathogenic (P1 and P2) or saprophytes (S1, S2) bacteria, were obtained and aligned by ClustalX software (version 2.1). Phylogenetic tree was built with ClustalW software (version 2.1).

### Statistical analysis

Prism 9 (GraphPad Software) was employed for all the statistical analysis of *in vivo* data.

Fisher’s exact test and analysis of variance (ANOVA) with Bonferroni’s multiple comparisons post-tests were applied to assess statistical differences between pairs of groups and multiple groups, respectively. Experiments for degradation of ECM substrates and C3b were carried out with three or four replicates. Data were presented as mean ± SD or SEM as shown in the figures. All data were assessed by SPSS 11.5 Software. A *P* value of <0.05 was considered significant.

## Acknowledgements

We would like to thank Matilde Costa Lima de Souza and Daniella dos Santos Courrol (Laboratory of Bacteriology, Instituto Butantan, São Paulo, Brazil), and Dr. Karukriti K. Ghosh (Yale School of Public Health, CT, US) for technical assistance. We also wanted to thank Dr. Mathieu Picardeau (Institut Pasteur, Paris, France) for providing the *Himar1* mutant of MPL36 (LIC10054) used in our studies.

---

**S1 Fig. Development and characterization of MPL36 mutants.** (A) Schematic representation of Himar1 transposon insertion positions in *L. interrogans* Manilae L495. The insertion sites of the transposon in the chromosome of the *mpl36* gene in strain Manilae WT, and the insertion site of the transposon containing the spectinomycin resistance cassette and *mpl36* gene for complementation are indicated. (B) Immunoblot analysis of Manilae WT, mutant Δ*mpl36* and complemented strain Δ*mpl36*^+^ using a rabbit polyclonal antibody against rMPL36. The visible band has a molecular weight of ∼40 kDa, which is in accordance with the predicted molecular weight of MPL36. The mutant strain lacks the expression of the protein. Molecular mass markers are shown on the left. (C) Growth curve analysis of Manilae WT, Δ*mpl36*, and Δ*mpl36*^+^ strains at 30°C. Bacteria were grown in EMJH medium without agitation, and the counting was performed using dark field microscopy. Results represent the average ± the standard deviation of three independent experiments.

**S2 Fig. Surface localization of MPL36 and immunogenicity in humans**. (A) Whole intact Manilae WT (WCP) was treated with Triton X-114 for phase partitioning of *Leptospira* membrane proteins. Immuno-blot analysis was conducted with detergent (D) and aqueous (A) phase using polyclonal rabbit antisera against LigA (outer membrane), GroEL (cytoplasmic), and MPL36 proteins. Antibodies to rMPL36 in human sera from individuals with confirmed severe leptospirosis were measured by an ELISA assay. Reactivity of the rMPL36 with acute and convalescent serum samples was tested for IgM (B) and IgG (C) levels separately. The dashed line represents the threshold calculated based on 2.5 SD of the average OD signal of sera from healthy US individuals used as control. Data show the mean absorbance value at 450 nm ± the standard deviation of all individuals tested.

**S3 Fig. Structural and phylogenetic analysis of MPL36 across species**. (A) Tertiary predicted model of the SPOR domain of MPL36 in *L. interrogans*, visualized by PyMOL, showing the exposed lysine residues in blue. (B) Alignment of tertiary predicted structures of SPOR domain from *L. interrogans* (P1-red), *L. fainei* (P2-pink), and *L. biflexa* (S1-green), visualized by PyMOL, showing the exposed lysine residues. (C) Dendrogram resulting from multiple alignments performed by ClustalW of SPOR domain of MPL36 with other similar sequences of all 69 *Leptospira* species separated by groups: P1 (red), P2 (pink), S1 (green), and S2 (blue), identified by BLASTp. (D) Alignment of the SPOR domain of MPL36 with other similar sequences of all 69 *Leptospira* species: P1 (red), P2 (pink), S1 (green), and S2 (blue). Alignment was done using the ClustalX software and similar amino acids have the same colors.

**S4 Fig**. **Assessment of ECM and complement C3b degradation by PLA bound to *Leptospira* strains**. Strains Manilae WT, Δ*mpl36*, Δ*mpl36*+ (10^8^ cells) were incubated with purified human PLG (10 µg). After washing, laminin (5 µg), vitronectin (2.5 µg) or C3b (1.5 µg) plus uPA (3 U) were added and incubated for up to 18 h. Leptospiral supernatants were collected and analyzed by western blot using anti-human vitronectin, laminin, or C3b (1:5,000) followed by peroxidase-conjugated secondary antibodies (1: 10,000).

